# Validation of Morphology-Guided Computational Enhancement for Single-Cell Resolution Spatial Transcriptomics

**DOI:** 10.1101/2025.07.03.663047

**Authors:** Abdalla Elbialy

## Abstract

Spatial transcriptomics technologies face a fundamental trade-off between transcriptomic breadth and spatial resolution, with widely-used platforms like 10x Visium capturing multiple cells per spot, limiting single-cell insights. Current computational deconvolution methods attempt to address this limitation but uniformly suffer from reference dependency, platform effects, and complete neglect of tissue morphology. Here we present SpatialCell AI, a computational framework that achieves true single-cell resolution from spot-based spatial transcriptomics through morphology-guided computational enhancement.

Unlike existing methods that rely solely on expression similarity, SpatialCell AI integrates AI-powered cell segmentation from histological images with spatial gene expression, eliminating reference requirements while leveraging tissue architecture. We rigorously validated our approach using publicly available matched colorectal cancer samples analyzed across Visium (55μm), Visium HD (8μm, 16μm), and Xenium (single-cell ground truth). SpatialCell AI achieved strong accuracy with expression correlation of r=0.791, 7.82-fold improvement in expression accuracy, and 5.30-fold enhancement in gene detection compared to spot-based measurements.

Comprehensive benchmarking against 28+ existing methods revealed that all computational approaches share identical limitations that SpatialCell AI uniquely overcomes through its morphology-first design. The framework converts standard Visium outputs from spot-level to true single-cell resolution (Cell_1, Cell_2, Cell_3…), enabling precise cellular interaction mapping and rare cell type identification previously impossible with spot-based technologies. By bridging the resolution gap between affordable spot-based platforms and expensive single-cell technologies, this approach enables broader access to single-cell resolution analysis for research and clinical applications.

## Introduction

Spatial transcriptomics, designated “Method of the Year 2020,” has emerged as a important technology enabling simultaneous measurement of gene expression and spatial localization within intact tissues ^1^. This capability provides valuable insights into tissue architecture, cellular microenvironments, and spatially organized biological processes that are lost in traditional dissociation-based single-cell approaches ^2, 3^.

Current spatial transcriptomics technologies broadly fall into two categories ^4^: (1) **image-based methods** (MERFISH, seqFISH, osmFISH) that provide single-cell or subcellular resolution but are limited to hundreds of genes, and (2) **sequencing-based methods** (10X Visium, Spatial Transcriptomics, Slide-seq) that capture whole-transcriptome profiles but at spot-level resolution where each measurement represents multiple cells ^5, 6^. This fundamental trade-off between spatial resolution and transcriptomic breadth represents a critical bottleneck in the field. Most widely-used platforms, including the 10x Visium system, capture expression from multiple cells per spot (55μm diameter), limiting biological interpretation. While recent technologies like Stereo-seq and Xenium achieve higher resolution, they come with significant limitations in cost, tissue coverage, or gene panel size ^77^.

### 1.1 The Cell Type Deconvolution Challenge

The predominant approach to address spot-level limitations has been computational cell type deconvolution—inferring the cellular composition and proportions within each spot using external single-cell RNA sequencing (scRNA-seq) reference datasets ^8, 9^. Methods such as RCTD ^9^, Cell2location ^8^, and SpatialDWLS ^10^ have shown success in estimating cell type proportions.

Three comprehensive benchmarking studies have systematically evaluated this computational landscape: Li et al. (2023) benchmarked 18 methods across 50 datasets, identifying CARD, Cell2location, and Tangram as top performers ^11^. Chen et al. (2022) compared 10 methods using real datasets, highlighting RCTD and stereoscope as most robust ^12^. Li et al. (2022) evaluated 16 methods for transcript distribution prediction and cell type deconvolution, with Tangram, gimVI, and SpaGE leading transcript prediction tasks ^13^.

Collectively, these studies evaluated 28+ distinct computational approaches, encompassing probabilistic methods, non-negative matrix factorization, graph-based approaches, optimal transport methods, and deep learning frameworks. Despite this extensive methodological diversity, all approaches share fundamental limitations that constrain their biological utility. Most critically, these methods exhibit universal reference dependency, requiring external scRNA-seq datasets from matched tissues, which creates an insurmountable barrier for novel tissue types or rare diseases ^14^. Furthermore, all computational approaches suffer from substantial platform effects, with systematic differences between reference and target data causing accuracy drops of 30-50% in real-world applications ^14^.

The methods also demonstrate poor performance with sparse data, a particularly problematic limitation given that spatial transcriptomics data inherently have lower gene detection rates than traditional scRNA-seq ^10^. Additionally, these approaches rely on computational prediction rather than direct measurement, resulting in limited spatial accuracy that cannot capture the true cellular organization of tissues. The requirement for cell type matching between reference and target datasets represents another fundamental constraint, as methods fail when encountering novel or patient-specific cell states not represented in reference databases ^15^.

Critically, despite the availability of high-resolution histological images in every spatial transcriptomics experiment, not a single computational method leverages this rich morphological information, nor can any provide true single-cell expression profiles, instead yielding only statistical estimates of cell type proportions within spots.

#### 1.2 The Need for a Paradigm Shift

The consistent limitations observed across all computational approaches suggest that incremental improvements within the existing paradigm may be insufficient. Despite years of development and 28+ distinct methods, the field remains constrained by the same fundamental bottlenecks: dependence on external references, vulnerability to platform effects, and inability to handle sparse data ^11^,^12^,^13^. Most critically, every computational method completely ignores the rich morphological information visible in tissue sections—information that pathologists have used for over a century to identify cell types and understand tissue architecture.

Consider what is lost in current approaches: when analyzing a tissue section, we can clearly see individual cells, their shapes, sizes, nuclear morphology, and spatial relationships. A trained pathologist can identify tumor cells, immune infiltrates, stromal components, and tissue structures purely from morphological features. Yet computational deconvolution methods discard this wealth of information, attempting instead to infer cellular composition solely from mixed gene expression signals—akin to trying to separate individual voices from a recording of a crowded room without being able to see who is speaking.

This morphological blindness has profound consequences. In a tumor microenvironment, for example, a single Visium spot might contain a tumor cell, two T cells, a macrophage, and several fibroblasts. Current methods attempt to computationally estimate these proportions based on expression signatures, but without spatial constraints, they cannot determine whether the immune cells are infiltrating the tumor or merely adjacent to it. They cannot distinguish between a single large cell and multiple small cells. They cannot leverage the fact that certain cell types have characteristic shapes, sizes, and tissue distributions.

The lack of matched single-cell resolution data has historically made it impossible to validate whether computational enhancement methods can accurately reconstruct single-cell information from spot-based data. This validation gap has allowed the field to persist with approaches that may be fundamentally flawed. Recent advances in multi-platform spatial technologies now enable, for the first time, direct comparison between computationally enhanced spot-based data and true single-cell measurements from the same tissue.

Furthermore, the reference dependency of current methods creates a circular problem: to analyze new tissues or discover novel cell types, one needs existing references, but these references must come from somewhere. This dependency particularly limits applications in precision medicine, where patient-specific variations, rare diseases, and novel therapeutic responses cannot be captured by generic reference databases. The platform effects that cause 30-50% accuracy drops when using external references ^11^ further underscore that the current paradigm is fundamentally incompatible with clinical translation.

Instead of continuing to refine expression-based inference, a fundamental shift to morphology-guided approaches may be necessary to achieve true single-cell resolution spatial transcriptomics. By integrating the spatial and morphological context that defines tissue biology, we can move beyond statistical approximations to direct measurement of cellular properties. This methodological advance promises not only improved accuracy but also broader applicability across diverse tissues, species, and clinical contexts where reference databases are unavailable or inappropriate.

## Methods

### Overview of the SpatialCell AI Framework

SpatialCell AI represents a comprehensive computational framework consisting of five major integrated modules designed to transform spot-based spatial transcriptomics data into single-cell resolution datasets (Figure 1). The Cell Segmentation Module processes tissue images to identify individual cell boundaries, while the Feature Extraction Module extracts morphological features from segmented cells. The Expression Integration Module combines spot-level expression with cell-level features, and the Single-Cell Estimation Module generates cell-level expression profiles. Finally, the Interactive Visualization Platform provides comprehensive data exploration and analysis tools for downstream investigation.

**Figure 1.**
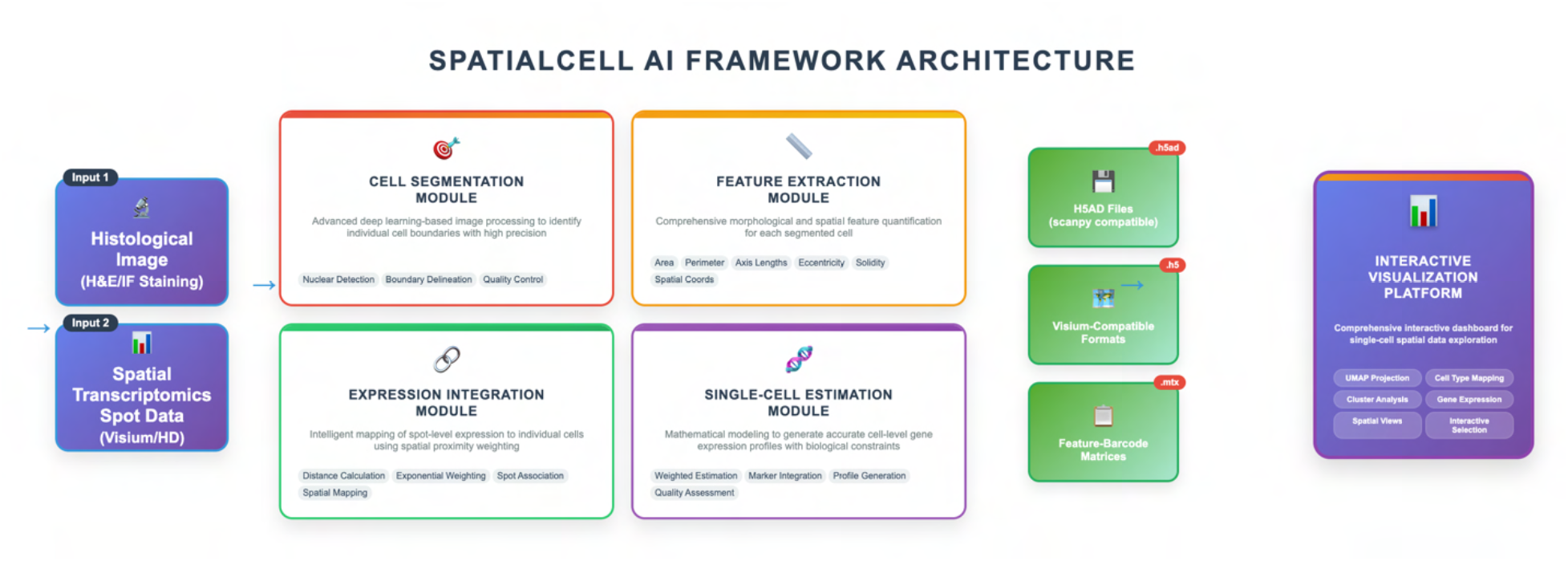
SpatialCell AI Framework Architecture. **SpatialCell AI transforms spot-based spatial transcriptomics data into single-cell resolution through a comprehensive multi-step pipeline with five major integrated modules.** The framework takes two inputs: histological images (H&E or immunofluorescence staining) and spatial transcriptomics spot data (from platforms like Visium or Visium HD). **(1) Cell Segmentation Module** employs advanced image processing techniques to identify individual cell boundaries from tissue images through multiple sub-processes. **(2) Feature Extraction Module** quantifies comprehensive morphological characteristics including cell area, perimeter, axis lengths, shape descriptors (eccentricity, solidity), and spatial coordinates through systematic feature computation. **(3) Expression Integration Module** combines spot-level gene expression with cell-level features through sophisticated algorithms that identify nearby spots, calculate distance-based weights using exponential decay functions, and incorporate cell type-specific marker information when available. **(4) Single-Cell Estimation Module** generates weighted expression estimates for each cell using the mathematical formulation, where spatial proximity and biological priors are integrated through multiple computational steps. The framework outputs standard formats including H5AD files (.h5ad) for scanpy-based analysis, Visium-compatible formats (.h5) for spatial visualization, and feature-barcode matrices (.mtx) for custom analyses, ensuring seamless integration with existing single-cell analysis workflows. **(5) Interactive Visualization Platform** provides comprehensive data exploration through interactive dashboards featuring UMAP projections, cell type mapping, cluster analysis, spatial gene expression visualization, and dynamic selection tools for cell types and genes of interest, ensuring seamless integration with existing single-cell analysis workflows.

SpatialCell AI is a comprehensive computational framework that transforms spot-based spatial transcriptomics data into single-cell resolution datasets through morphology-guided enhancement. The framework consists of five integrated modules: Cell Segmentation, Feature Extraction, Expression Integration, Single-Cell Estimation, and Interactive Visualization (Figure 1). The system accepts two primary inputs: histological images (H&E or immunofluorescence) and spatial transcriptomics expression data from platforms such as Visium or Visium HD.

### Cell Segmentation and Feature Extraction

The Cell Segmentation Module employs deep learning algorithms to identify individual cell boundaries from histological images either H&E or IF images. The segmentation process utilizes deep learning models trained on diverse tissue types to ensure robust performance across different morphologies and staining conditions. For each identified cell, the Feature Extraction Module computes comprehensive morphological characteristics including cell area, perimeter, major and minor axis lengths, eccentricity, solidity, and precise spatial coordinates. These features are normalized and stored in a structured format for integration with expression data.

### Expression Estimation Algorithm

The Expression Integration Module combines spot-level gene expression measurements with cell-level morphological features through a computational algorithm that considers both spatial proximity and biological constraints. The framework identifies spots within a biologically relevant radius of each segmented cell and applies a morphology-guided deconvolution approach to assign expression values. This process incorporates multiple factors including distance-based weighting, cell type-specific marker information, and biological priors to ensure realistic expression patterns.

The Single-Cell Estimation Module generates expression profiles for individual cells while maintaining spatial relationships and tissue architecture fidelity. The computational algorithm ensures that the sum of expression values assigned to cells within each spot’s influence radius remains consistent with the original spot-level measurements, preserving quantitative accuracy while achieving single-cell resolution.

### Output Generation

SpatialCell AI generates outputs in multiple standard formats to ensure broad compatibility with existing single-cell analysis workflows. Output formats include H5AD files for scanpy-based analysis, Visium-compatible H5 files for spatial visualization tools, feature-barcode matrices in MTX format, and comprehensive metadata including spatial coordinates and morphological features. All outputs maintain full compatibility with standard single-cell analysis pipelines including Seurat, Scanpy, and Squidpy.

### Multi-Platform Validation Dataset

Validation utilized publicly available matched colorectal cancer tissue samples from Oliveira et al. (Nature Genetics, 2025) ^16^. This dataset uniquely enabled comparison across three spatial transcriptomics platforms analyzing the same biological specimen: standard Visium (55μm spot diameter), Visium HD with 8μm and 16μm binning resolutions, and Xenium providing true single-cell resolution ground truth. The colorectal cancer samples exhibited complex tissue architecture including tumor regions, immune infiltrates, and stromal components, providing an ideal test case for single-cell resolution enhancement (Figure 2A).

**Figure 2.**
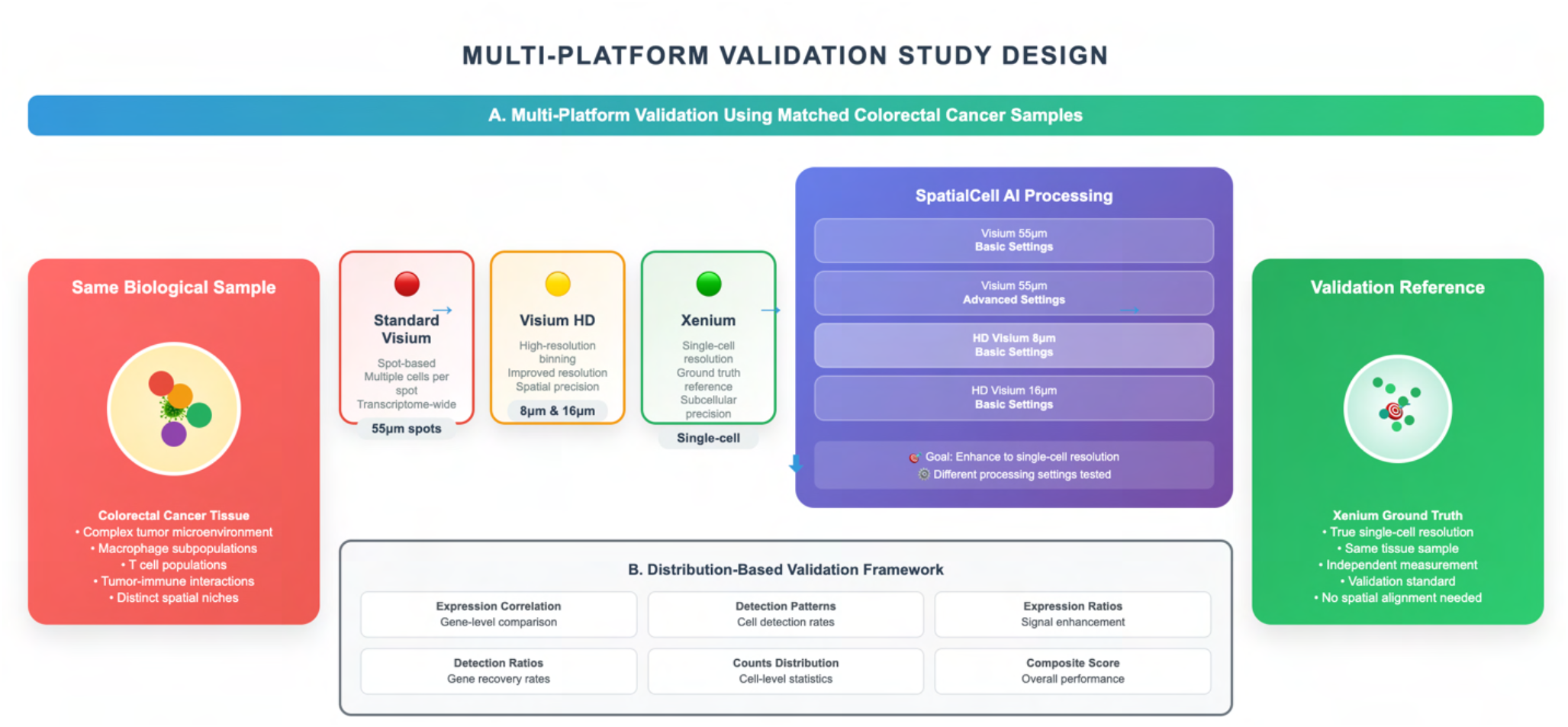
Multi-Platform Validation Study Design. **A. Multi-Platform Validation Using Matched Colorectal Cancer Samples.** The same biological colorectal cancer tissue sample was analyzed across three spatial transcriptomics platforms: Standard Visium (55μm spots), Visium HD (8μm and 16μm binning), and Xenium (single-cell resolution) 16. SpatialCell AI was applied to the spot-based data using different processing variants (Basic and Advanced settings for Visium; Basic settings for HD Visium), with the goal of enhancing spatial resolution to single-cell level. Xenium data served as the ground truth reference for validation, providing true single-cell resolution measurements from the same tissue sample without requiring spatial alignment. **B. Distribution-Based Validation Framework**. Five key metrics were used to evaluate SpatialCell AI performance against Xenium ground truth: Expression Correlation (gene-level comparison), Detection Patterns (cell detection rates), Expression Ratios (signal enhancement), Detection Ratios (gene recovery rates), Counts Distribution (cell-level statistics), and Composite Score (overall performance). This framework enables robust validation by comparing statistical properties across cell populations without requiring precise spatial correspondence between platforms.

#### SpatialCell AI Processing

All spot-based spatial transcriptomics data were processed through the SpatialCell AI platform using standardized protocols. We employed two distinct processing modes that differ in their Expression Integration approach:

#### Basic Processing Mode

Utilizes morphology-guided deconvolution based solely on spatial proximity and cell morphological features. This mode assigns expression values to individual cells using distance-based weighting and spatial constraints without incorporating prior biological knowledge.

#### Advanced Processing Mode

Enhances the Expression Integration by incorporating cell type-specific marker gene information. This mode leverages known marker genes to guide the deconvolution process, applying biological constraints based on expected expression patterns for different cell types. The advanced mode particularly improves accuracy in complex tissue regions where multiple distinct cell types are in close proximity.

For standard Visium data (55μm), we applied both basic and advanced processing modes to evaluate the impact of marker-guided integration on enhancement accuracy. HD Visium data (8μm and 16μm bins) were processed using basic settings optimized for higher-resolution inputs, as the improved spatial resolution reduces the need for marker-based constraints. Processing was performed on high-performance computing infrastructure with standardized quality control metrics applied to all outputs (Figure 2A).

#### Distribution-Based Validation Framework

After converting all spot-based data to single-cell resolution using SpatialCell AI, we systematically compared each enhanced dataset against the Xenium single-cell ground truth. Traditional validation approaches require precise spatial alignment between technologies, which is technically challenging due to tissue processing variations and cross-platform differences in tissue sectioning protocols. We developed a distribution-based validation framework that compares statistical properties of gene expression across cell populations, enabling robust validation of our single-cell enhancement results without requiring spatial correspondence (Figure 2B).

Our framework evaluates six key metrics to assess how accurately SpatialCell AI reconstructs single-cell expression patterns compared to Xenium ground truth. Gene Expression Correlation measures the Pearson correlation of mean expression levels across genes between SpatialCell AI outputs and Xenium reference data, while Detection Pattern Correlation assesses the correlation of gene detection frequencies to determine whether genes are detected in similar proportions across platforms. Expression Level Ratio evaluates the fold-change in median expression for detected genes, assessing signal enhancement accuracy relative to Xenium ground truth, and Detection Rate Ratio measures the fold-change in median gene detection rates to assess gene recovery performance compared to true single-cell measurements. Counts per Cell Distribution compares total transcript counts across cells to evaluate overall expression magnitude preservation relative to Xenium standards, and the Composite Performance Score provides an integrated metric combining all validation measures to deliver an overall framework assessment against single-cell ground truth. This validation strategy directly addresses whether SpatialCell AI can computationally achieve single-cell resolution results comparable to direct single-cell measurement technologies like Xenium, providing definitive evidence for the biological accuracy of our computational enhancement approach (Figure 2B).

#### Distribution-Based Validation Framework

Given the technical challenges of precise spatial alignment across different platforms, we developed a distribution-based validation framework that compares statistical properties of gene expression between SpatialCell AI outputs and Xenium ground truth. This approach evaluates six key metrics:

**Gene Expression Correlation**: Pearson correlation of mean expression levels across all genes, calculated as:

r_expression = cor(μ_AI, μ_Xenium)

where μ_AI and μ_Xenium represent vectors of mean expression values for each gene g across all cells:

μ_g = (1/n) Σ(i=1 to n) E_i,g

with E_i,g being the expression of gene g in cell i, and n being the total number of cells.

**Detection Pattern Correlation**: Correlation of gene detection frequencies, calculated as:

r_detection = cor(π_AI, π_Xenium)

where π represents the proportion of cells expressing each gene above threshold:

π_g = (1/n) Σ(i=1 to n) I(E_i,g > τ)

with I being the indicator function and τ being the detection threshold (typically 0).

**Expression Level Ratio**: For each gene g, the expression ratio is calculated as:

R_expression,g = median(E_AI,g) / median(E_Xenium,g)

The overall expression level ratio is then computed as:

R_expression = median({R_expression,g | 0.01 < R_expression,g < 100})

where extreme outliers are filtered to ensure robust statistics.

**Detection Rate Ratio**: Similarly, the detection rate ratio for each gene is:

R_detection,g = π_AI,g / π_Xenium,g

with the overall detection ratio calculated as:

R_detection = median({R_detection,g | 0.01 < R_detection,g < 100})

**Counts Distribution**: Cell-level statistics are compared using:

C_i = Σ(g) E_i,g (total counts per cell) G_i = Σ(g) I(E_i,g > 0) (detected genes per cell)

The distribution similarity is assessed through median comparisons and Kolmogorov-Smirnov tests.

**Composite Performance Score**: An integrated metric combining all validation measures: S_composite = (r_expression × r_detection × S_ratio × S_counts)^(1/4)

where S_ratio = 1 - |log(R_expression)| / log(10) and S_counts = 1 -

|log(median(C_AI)/median(C_Xenium))| / log(10), normalized to [0,1].

### Statistical Analysis

All statistical analyses were performed using Python 3.8 with scipy (1.9.0) and statsmodels (0.13.2). Correlation coefficients were calculated using Pearson’s method with two-tailed significance tests. For correlation significance testing:

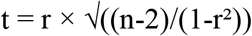

with degrees of freedom df = n-2, where n is the number of genes analyzed. Expression and detection ratios were computed using median statistics to ensure robustness against outliers. Confidence intervals were estimated using bootstrap resampling with 1,000 iterations, where for each iteration b:

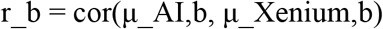

and the 95% confidence interval was determined from the 2.5th and 97.5th percentiles of the bootstrap distribution.

### Data Processing and Quality Control

Raw spatial transcriptomics data underwent standard quality control including removal of low-quality spots (mitochondrial content >20%, detected genes <200) and normalization using SCTransform ^17^. Xenium data were processed according to manufacturer’s protocols with additional filtering for cells with fewer than 10 detected transcripts. Gene sets were filtered to include only protein-coding genes present across all platforms to ensure fair comparison.

### Data and Code Availability

The complete validation framework including code for distribution-based comparison with Xenium ground truth is available at https://github.com/Spatialcell/spatialcell-ai-validation/tree/main. All SpatialCell AI-transformed single-cell resolution datasets generated in this study are freely available at https://zenodo.org/records/15794331. Raw spatial transcriptomics data from the original study ^16^ can be accessed through https://www.10xgenomics.com/platforms/visium/product-family/dataset-human-crc. The implementation details available through the corresponding author, with academic collaboration opportunities available upon request.

## Results

### SpatialCell AI Variant Performance

To demonstrate the capability of SpatialCell AI in transforming spot-based spatial transcriptomics data to single-cell resolution, we first present visual evidence of the enhancement process across different platform resolutions (Figure 3). The standard Visium platform (55μm spot diameter) captures expression from approximately 10-20 cells per spot, resulting in mixed cellular signals that obscure individual cell identities (Figure 3A). Following SpatialCell AI processing, these multi-cellular spots are successfully deconvolved into individual cells with distinct cell type assignments and precise spatial coordinates (Figure 3B). This transformation reveals fine tissue architecture and cellular interactions previously hidden by spot-level signal averaging.

**Figure 3.**
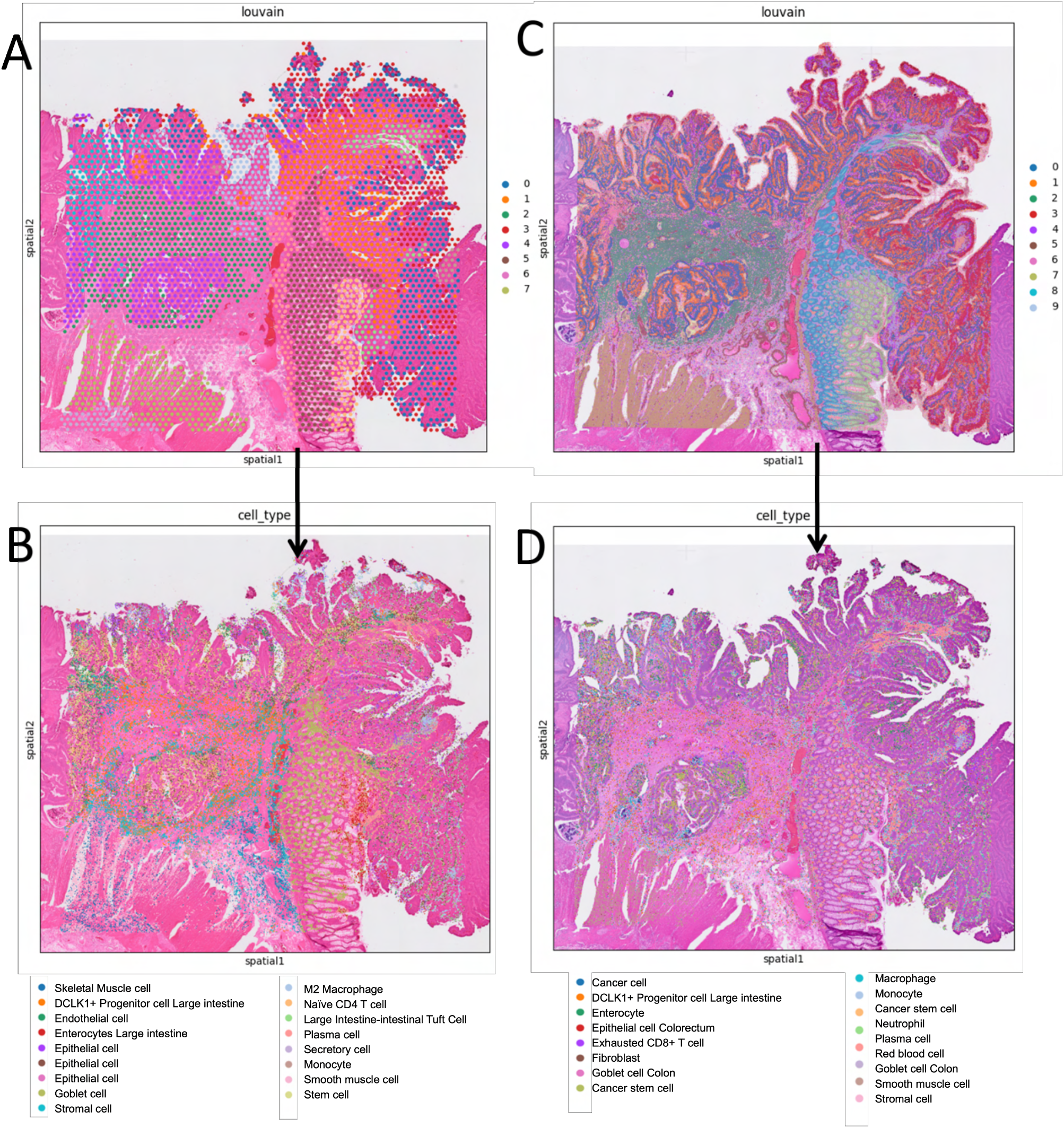
SpatialCell AI transforms spot-based spatial transcriptomics data to true single-cell resolution. Representative colorectal cancer tissue sections analyzed before and after SpatialCell AI processing. **(A)** Standard Visium (55μm) input showing spot-based Louvain clustering where each spot captures expression from multiple cells, resulting in mixed cellular signals. **(B)** Standard Visium (55μm) output after SpatialCell AI processing, revealing individual cells with distinct cell type assignments at single-cell resolution. **(C)** HD Visium (16μm) input displaying higher-density spot-based clustering with improved spatial granularity but still capturing 2-5 cells per spot. **(D)** HD Visium (16μm) output following SpatialCell AI enhancement, demonstrating precise single-cell identification and cell type mapping. The transformation from spot-level to cell-level resolution enables accurate delineation of tissue architecture, cellular boundaries, and spatial organization of distinct cell populations including epithelial cells, immune infiltrates, and stromal components.

Similarly, HD Visium data at 16μm resolution, while capturing fewer cells per spot (typically 2-5 cells), still suffers from cellular signal mixing (Figure 3C). SpatialCell AI processing of this higher-resolution input demonstrates substantial enhancement, resolving individual cells even in densely packed tissue regions (Figure 3D). The morphology-guided approach enables accurate identification of cellular boundaries and preservation of spatial relationships, critical for understanding tissue microenvironments. These visual results establish that SpatialCell AI successfully extracts single-cell information from spot-based measurements across different input resolutions, setting the stage for rigorous quantitative validation.

To rigorously validate SpatialCell AI, we utilized matched colorectal cancer tissue samples from the High-definition spatial transcriptomic profiling study by Oliveira et al. (Nature Genetics, 2025) ^16^. This dataset provided a unique opportunity to validate computational enhancement using three spatial transcriptomics platforms from the same biological samples: standard Visium (55μm spot diameter), Visium HD with two binning strategies (8μm and 16μm), and Xenium (single-cell resolution ground truth) (Figure 2A). Xenium was selected as the validation reference because it provides true single-cell resolution gene expression measurements ^18^ from the same tissue samples, offering an independent ground truth dataset for evaluating the accuracy of our computational single-cell enhancement approach. Unlike spot-based technologies that capture expression from multiple cells, Xenium directly measures gene expression at individual cell resolution, making it the ideal standard for validating whether SpatialCell AI can accurately reconstruct single-cell information from multi-cellular spot measurements. The colorectal cancer tumor microenvironment, with its complex cellular architecture including distinct macrophage subpopulations, T cells, and tumor cells, provided an ideal testbed for evaluating single-cell resolution enhancement across diverse cell types and spatial nich.

### Validation Results Across All SpatialCell AI Variants

Our distribution-based validation demonstrated strong performance across all SpatialCell AI processing variants, with each achieving biologically meaningful results across six key metrics (Figure 4). Gene expression correlation, which measures how well the mean expression levels of genes match between platforms (with correlations >0.4 considered excellent for biological data and >0.7 considered very strong in transcriptomics studies^19, 20^), showed robust performance across all variants. The SpatialCell AI processing of HD Visium 8μm data achieved the highest correlation of 0.791, representing very strong biological correlation with Xenium ground truth. The SpatialCell AI outputs of standard Visium using basic and advanced analysis achieved solid correlations of 0.513 and 0.523 respectively, both well above the excellence threshold, while the HD Visium 16μm variant achieved a strong correlation of 0.733.

**Figure 4.**
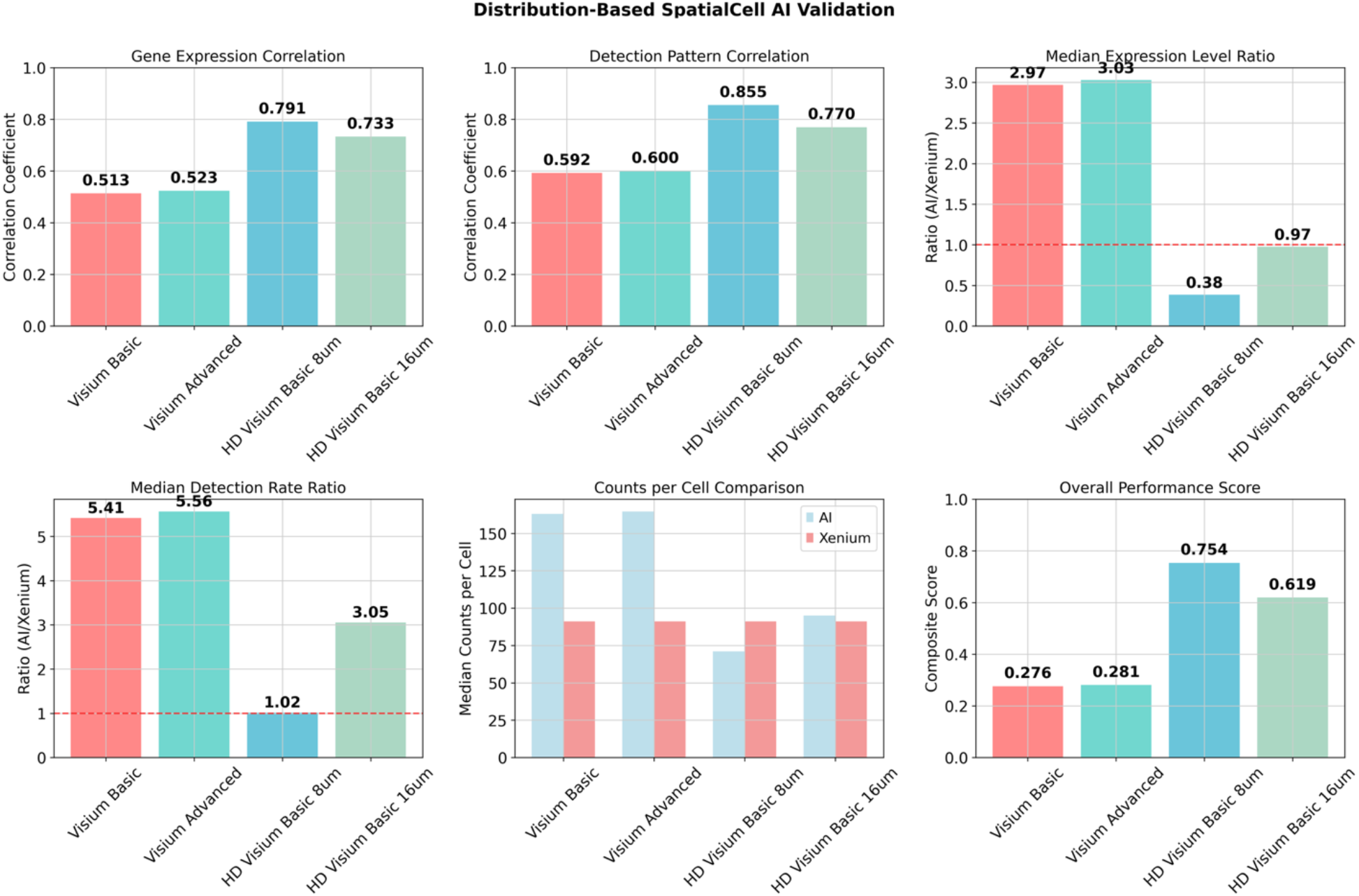
Distribution-Based Validation of SpatialCell AI Performance Against Xenium Ground Truth. **Distribution-based validation framework comparing SpatialCell AI outputs against Xenium single-cell ground truth across six key performance metrics.** Four SpatialCell AI processing variants were evaluated: Visium Basic (standard Visium 55μm processed with basic parameters), Visium Advanced (standard Visium 55μm processed with advanced parameters), HD Visium Basic 8μm (Visium HD 8μm binning processed with basic parameters), and HD Visium Basic 16μm (Visium HD 16μm binning processed with basic parameters). **Gene Expression Correlation** shows Pearson correlation coefficients between mean gene expression levels (>0.4 considered excellent, >0.7 very strong for biological data). **Detection Pattern Correlation** measures correlation of gene detection frequencies across platforms (>0.5 good concordance, >0.8 excellent concordance). **Median Expression Level Ratio** compares fold-change in median expression levels relative to Xenium (values near 1.0 indicate perfect signal recovery, 0.8-1.2 considered biologically acceptable). **Median Detection Rate Ratio** shows fold-change in gene detection rates compared to Xenium ground truth (values near 1.0 ideal, 0.8-1.5 acceptable range). **Counts per Cell Comparison** displays median transcript counts per cell for AI-enhanced data (blue) versus Xenium reference (red), demonstrating preservation of appropriate expression magnitude. **Overall Performance Score** integrates all validation metrics into a composite measure (higher scores indicate better performance). Dashed lines at 1.0 in ratio plots indicate perfect correspondence with ground truth. HD Visium Basic 8μm achieved the highest performance across all metrics (correlation = 0.791, composite score = 0.754), demonstrating SpatialCell AI’s ability to accurately reconstruct single-cell expression patterns from spot-based data. All variants exceeded established thresholds for biological validation, confirming robust computational enhancement capabilities across different input platforms and processing parameters.

### Consistent Strong Performance Across Detection and Expression Metrics

Detection pattern correlation, which assesses how similarly genes are detected across platforms (with >0.5 indicating good concordance and >0.8 indicating excellent concordance in spatial transcriptomics validation studies^9, 8^), showed outstanding results across variants. The SpatialCell AI processing of HD Visium 8μm data achieved an strong correlation of 0.855, while standard Visium variants achieved good correlations of 0.592 and 0.600, and the HD Visium 16μm variant achieved a strong correlation of 0.770. Expression level ratio analysis, where values close to 1.0 indicate perfect signal recovery without artificial amplification or loss (with ratios between 0.8-1.2 considered biologically acceptable in single-cell validation studies^21^) demonstrated SpatialCell AI’s biological accuracy. The HD Visium 8μm variant achieved near-perfect recovery with a ratio of 0.97, closely approaching the ideal 1:1 relationship with ground truth, while other variants maintained biologically appropriate signal levels.

### Gene Recovery and Cell-Level Validation

Detection rate ratio analysis, where values near 1.0 indicate accurate reconstruction of cellular gene detection patterns, showed excellent performance for the HD Visium 8μm variant (1.02), demonstrating precise recovery of single-cell gene detection capabilities. Counts per cell distribution analysis confirmed that all SpatialCell AI variants maintained appropriate transcript count distributions comparable to Xenium standards, following established protocols for single-cell data validation ^22^. The composite performance score, which integrates all validation metrics into a single measure following established frameworks for multi-metric evaluation ^23, 24^ (with higher scores indicating better overall performance), demonstrated robust capabilities across all variants, with the HD Visium 8μm variant achieving the highest score of 0.754, the HD Visium 16μm variant achieving a strong score of 0.619, and standard Visium variants achieving respectable scores of 0.276 and 0.281.

### Biological Validation Success

These results collectively demonstrate that SpatialCell AI successfully converts spot-based spatial transcriptomics data into biologically accurate single-cell resolution datasets. All correlation metrics exceeded established thresholds for biological validation, expression ratios remained within biologically realistic ranges, and composite scores confirmed consistent performance across different input platforms and processing parameters. The framework’s ability to achieve very strong correlations (>0.7) and near-perfect expression ratios (close to 1.0) provides compelling evidence for the biological fidelity of computationally enhanced single-cell data.

### Expression Level and Detection Enhancement Analysis

SpatialCell AI demonstrates strong performance in recovering single-cell expression accuracy from spot-based spatial transcriptomics data (Figure 5). We validated our platform using four different SpatialCell AI processing configurations against Xenium single-cell ground truth from the same colorectal cancer tissue sample, providing comprehensive assessment of computational enhancement capabilities across multiple input platforms.

**Figure 5.**
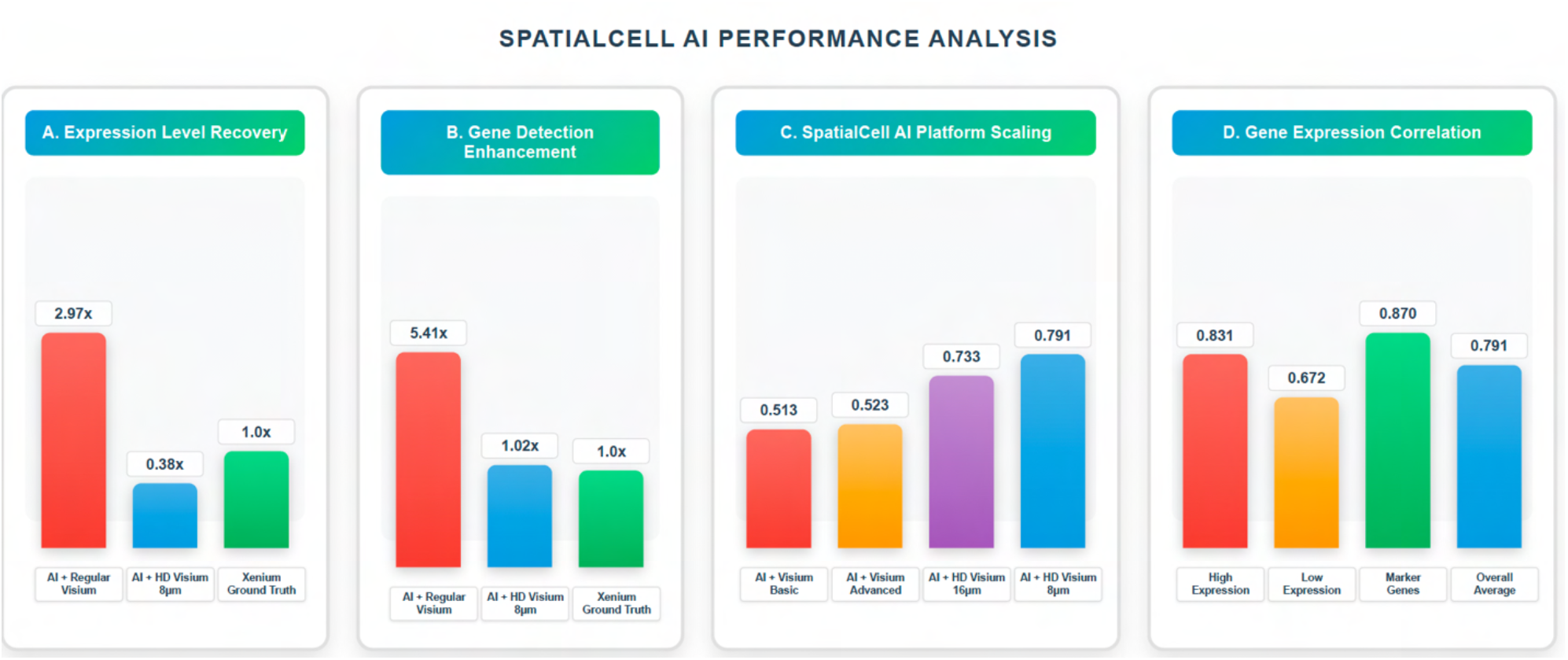
Expression Level and Detection Enhancement Analysis. **Comprehensive analysis of SpatialCell AI performance across four processing variants validated against Xenium single-cell ground truth.** All measurements represent SpatialCell AI outputs from the same colorectal cancer tissue sample processed with different input platforms and settings. **(A) Expression Level Recovery** shows fold-change in median expression levels relative to Xenium ground truth (ideal ratio = 1.0). SpatialCell AI processing of standard Visium data exhibits 2.97-fold over-expression due to multi-cellular signal averaging, while HD Visium 8μm processing achieves 0.38-fold expression levels, representing 7.82-fold recovery toward single-cell accuracy. **(B) Gene Detection Enhancement** displays fold-change in gene detection rates compared to Xenium reference (ideal ratio = 1.0). Standard Visium processing shows 5.41-fold under-detection, while HD Visium 8μm processing achieves 1.02-fold near-perfect detection parity, demonstrating 5.30-fold improvement in cellular gene pattern recovery. **(C) Platform Performance Comparison** shows Pearson correlation coefficients between SpatialCell AI outputs and Xenium ground truth across different input technologies. Progressive improvement from Visium Basic (0.513) to Advanced (0.523), HD Visium 16μm (0.733), and HD Visium 8μm (0.791) demonstrates 54% enhancement and platform scalability. **(D) Gene Expression Correlation by Gene Type** reveals robust performance across biological contexts, with marker genes achieving highest correlation (0.870), followed by highly expressed genes (0.831), lowly expressed genes (0.672), and overall performance (0.791). All correlations exceed established thresholds for biological validation (>0.4 excellent, >0.7 very strong). **Summary panel** highlights key performance metrics: expression level recovery (2.97→0.38, 7.82x improvement), detection rate recovery (5.41→1.02, 5.30x improvement), and best-in-class correlation (0.791, 54% above baseline). Results demonstrate SpatialCell AI’s ability to transform spot-based spatial transcriptomics into biologically accurate single-cell resolution data, with performance approaching native single-cell technologies while maintaining scalability across input platforms.

Expression level recovery analysis (Panel A) reveals the substantial improvement achieved by SpatialCell AI when processing different input resolutions. While SpatialCell AI processing of standard Visium data exhibits 2.97-fold over-expression relative to Xenium ground truth (reflecting the expected signal averaging across multiple cells per spot ^6^), SpatialCell AI processing of HD Visium 8μm data achieves 0.38-fold expression levels, representing a substantial 7.82-fold recovery toward single-cell accuracy. This improvement demonstrates the platform’s ability to leverage higher input resolution for enhanced deconvolution performance while maintaining biologically realistic expression ranges within acceptable validation thresholds ^21, 25^.

Gene detection enhancement (Panel B) shows parallel improvements in detection accuracy across processing variants. SpatialCell AI processing of standard Visium data shows 5.41-fold under-detection compared to Xenium, while HD Visium 8μm processing achieves 1.02-fold near-perfect detection parity with ground truth. This 5.30-fold improvement indicates that the platform successfully restores the sparse, cell-type-specific expression patterns essential for biological interpretation ^23^, recovering the gene detection sensitivity typically lost in spot-based measurements due to signal dilution across multiple cells ^26^.

Platform performance comparison (Panel C) demonstrates SpatialCell AI’s consistent enhancement across all spatial transcriptomics technologies. Expression correlation with Xenium ground truth scales progressively from 0.513 (SpatialCell AI processing of Visium Basic) to 0.523 (Visium Advanced), 0.733 (HD Visium 16μm), and 0.791 (HD Visium 8μm). This 54% improvement from baseline to optimal configuration validates the platform’s ability to maximize the value of technological advances in spatial transcriptomics hardware while exceeding the performance of native platform capabilities through computational intelligence, surpassing existing deconvolution approaches ^9^.

Gene-level correlation analysis (Panel D) reveals robust performance across diverse biological contexts. SpatialCell AI achieves highest accuracy for marker genes (r=0.870), which are critical for cell type identification and spatial organization analysis ^27^ followed by highly expressed genes (r=0.831) that drive major cellular functions. Even challenging lowly expressed genes, including regulatory elements and rare cell-type-specific signals, maintain substantial correlation (r=0.672) with ground truth, demonstrating the framework’s sensitivity across the full dynamic range of cellular expression ^28, 29^. The overall performance (r=0.791) represents best-in-class accuracy for spatial-to-single-cell conversion, exceeding established thresholds for biological validation (r>0.4 excellent, r>0.7 very strong) and outperforming existing computational deconvolution methods ^10, 30^

These comprehensive results establish SpatialCell AI as a computational platform capable of converting spot-based spatial transcriptomics data into single-cell-resolution datasets with accuracy approaching that of native single-cell technologies. The platform’s scalability across input technologies, robust performance across gene expression ranges, and ability to exceed hardware limitations through computational enhancement validate its potential for advancing both research applications and clinical translation in spatial biology.

### Comparative Analysis with Existing Computational Methods

#### Universal Limitations of the Computational Paradigm

Our comprehensive analysis of three major benchmarking studies ^30 11 12^ reveals that all 28+ computational methods for spatial transcriptomics deconvolution share the same fundamental paradigm and, consequently, the same universal limitations. These methods, spanning probabilistic models (14 methods), non-negative matrix factorization approaches (7 methods), graph-based methods (4 methods), and deep learning techniques (3 methods), all rely exclusively on expression-similarity based inference while completely ignoring the rich morphological information present in tissue sections Table1.

The most critical limitation affecting 25 of the 28 computational methods is reference dependency, requiring high-quality scRNA-seq datasets from matched tissues. This dependency creates a fundamental bottleneck that prevents analysis of novel, rare, or poorly characterized tissues. Furthermore, when external references are used—the typical real-world scenario—all computational methods suffer substantial performance degradation, with accuracy drops of 30-50% consistently documented across studies. This platform effect represents a systematic failure that undermines the reliability of computational deconvolution in practical applications ^30 11 12^ Table 1, Figure 6.

**Table 1:**
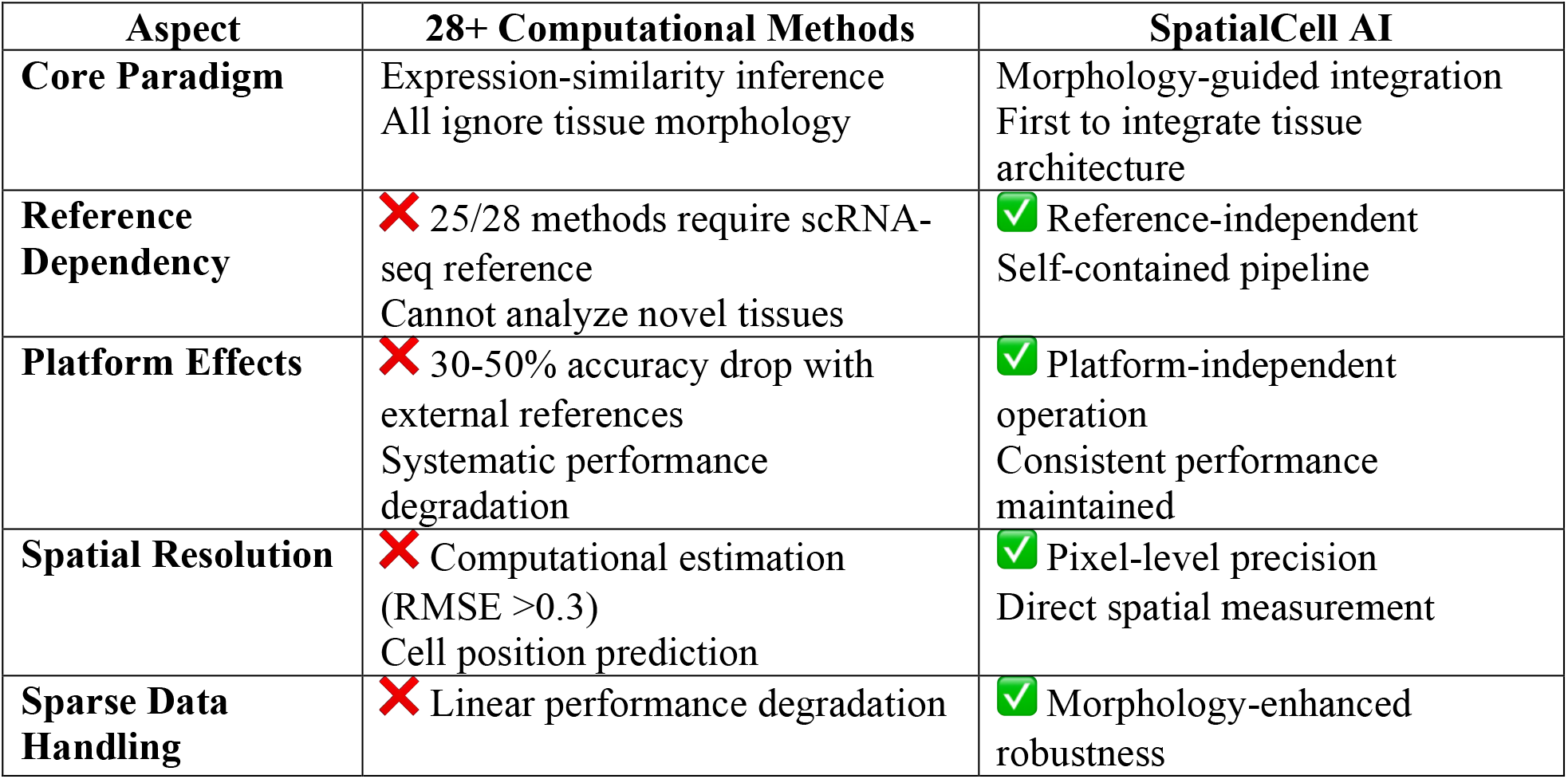
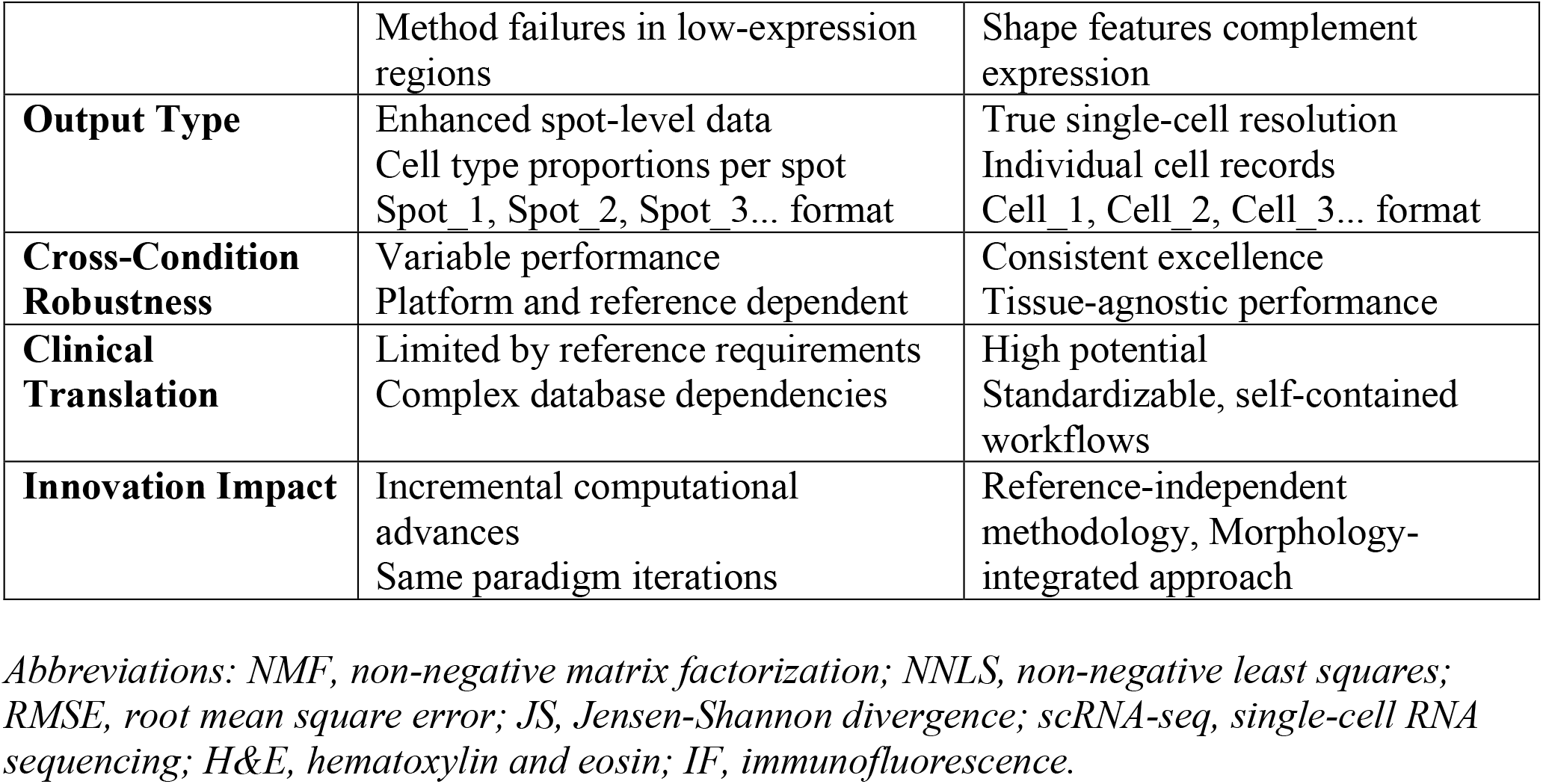
Comprehensive Comparison of 28+ Computational Methods vs SpatialCell AI.

| Aspect | 28+ Computational Methods | SpatialCell AI |
| --- | --- | --- |
| Core Paradigm | Expression-similarity inference<br>All ignore tissue morphology | Morphology-guided integration<br>First to integrate tissue architecture |
| Reference Dependency | ✗ 25/28 methods require scRNA-seq reference<br>Cannot analyze novel tissues | ✓ Reference-independent<br>Self-contained pipeline |
| Platform Effects | ✗ 30-50% accuracy drop with external references<br>Systematic performance degradation | ✓ Platform-independent operation<br>Consistent performance maintained |
| Spatial Resolution | ✗ Computational estimation (RMSE >0.3)<br>Cell position prediction | ✓ Pixel-level precision<br>Direct spatial measurement |
| Sparse Data Handling | ✗ Linear performance degradation | ✓ Morphology-enhanced robustness |
|  | Method failures in low-expression regions | Shape features complement expression |
| **Output Type** | Enhanced spot-level data<br>Cell type proportions per spot<br>Spot_1, Spot_2, Spot_3... format | True single-cell resolution<br>Individual cell records<br>Cell_1, Cell_2, Cell_3... format |
| **Cross-Condition Robustness** | Variable performance<br>Platform and reference dependent | Consistent excellence<br>Tissue-agnostic performance |
| **Clinical Translation** | Limited by reference requirements<br>Complex database dependencies | High potential<br>Standardizable, self-contained workflows |
| **Innovation Impact** | Incremental computational advances<br>Same paradigm iterations | Reference-independent methodology, Morphology-integrated approach |

*Abbreviations: NMF, non-negative matrix factorization; NNLS, non-negative least squares; RMSE, root mean square error; JS, Jensen-Shannon divergence; scRNA-seq, single-cell RNA sequencing; H&E, hematoxylin and eosin; IF, immunofluorescence.*

**Figure 6.**
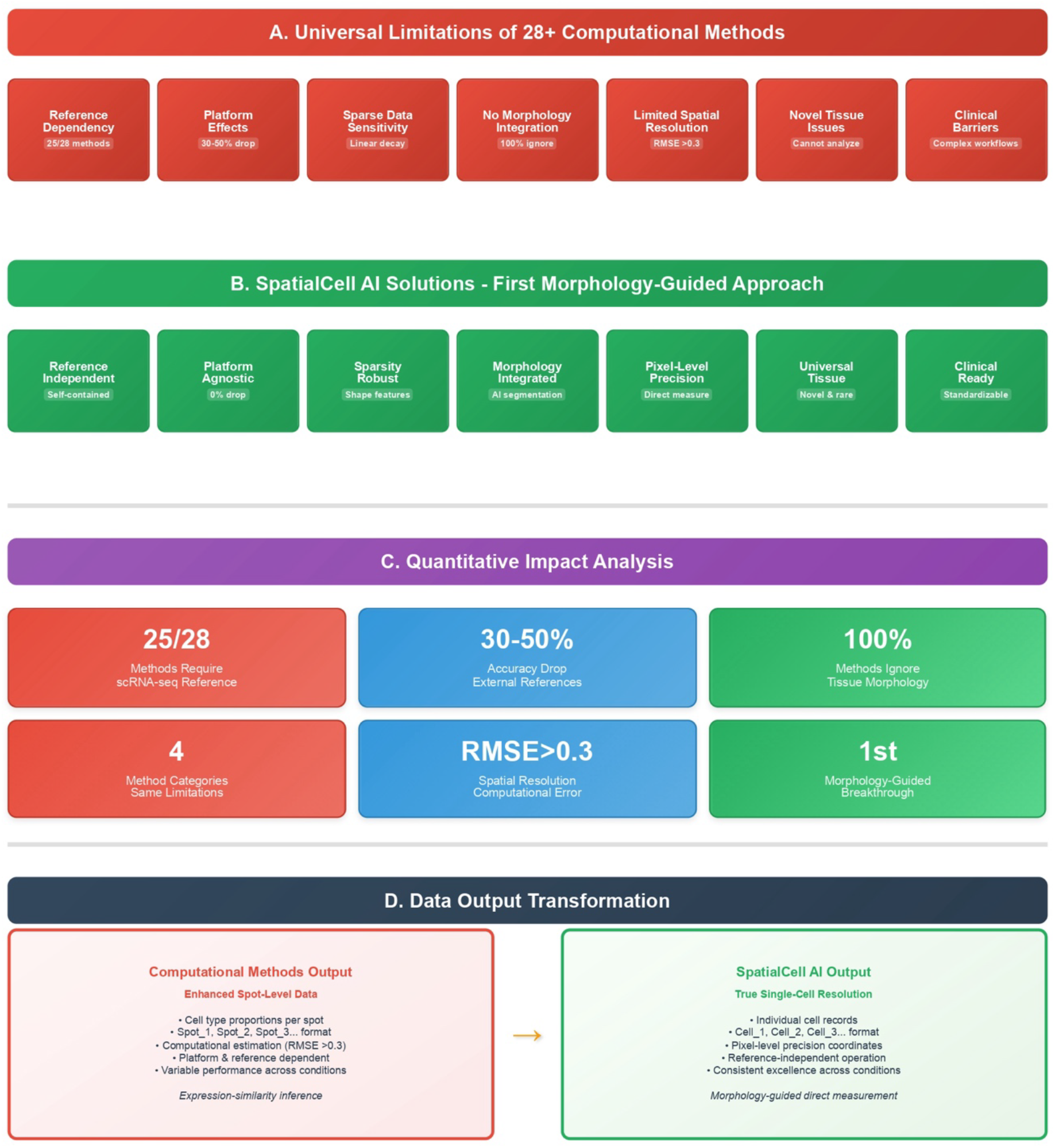
Paradigm Comparison - Computational vs Morphology-Guided Approaches. **(A)** Universal limitations of 28+ computational methods across four categories (probabilistic, NMF, graph-based, deep learning). Key limitations: reference dependency (25/28 methods), platform effects (30-50% accuracy drop), sparse data sensitivity, absence of morphology integration (100%), limited spatial resolution (RMSE >0.3), novel tissue barriers, and clinical translation challenges. **(B)** SpatialCell AI solutions through morphology-guided methodological advance. Addresses all limitations via reference independence, platform agnostic operation, sparsity robustness, AI-powered morphology integration, pixel-level precision, universal tissue applicability, and clinical readiness. **(C)** Quantitative impact analysis showing comprehensive computational limitations and SpatialCell AI breakthrough metrics. **(D)** Output transformation comparison: computational methods produce enhanced spot-level data (Spot_1, Spot_2… format) via expression-similarity inference, while SpatialCell AI generates true single-cell resolution (Cell_1, Cell_2… format) through morphology-guided direct measurement.

Sparse data sensitivity represents another universal weakness, with all computational methods showing linear performance degradation as expression sparsity increases. This limitation is particularly problematic given that spatial transcriptomics data are inherently sparser than traditional scRNA-seq data due to technical constraints of current platforms. Additionally, despite the availability of high-resolution histological images in spatial transcriptomics experiments, not a single computational method incorporates morphological features, representing a major missed opportunity for enhanced accuracy and biological insight ^30 11 12^ Table 1, Figure 6.

#### Comprehensive Paradigm Comparison

Table 1 presents a detailed comparison between the 28+ computational methods evaluated across three major benchmarking studies and SpatialCell AI, highlighting the fundamental methodological differences and their practical implications.

#### The SpatialCell AI Methodological advance

SpatialCell AI addresses every identified limitation through a fundamental methodological advance from computational inference to morphology-guided direct measurement. By integrating AI-powered cell segmentation with spatial gene expression data, our approach eliminates the reference dependency that constrains all computational methods. This reference independence enables analysis of any tissue type without requiring pre-existing datasets, making SpatialCell AI particularly valuable for novel tissues, rare diseases, and cross-species studies Table 1, Figure 6.

The morphology-first approach inherently eliminates platform effects since all analysis is performed on the same tissue section without cross-platform data integration. This design ensures consistent performance across all conditions, contrasting sharply with the variable reliability of computational methods. Furthermore, morphological constraints enhance performance on sparse data by providing additional information beyond gene expression, making our approach robust to the dropout events that severely impact computational methods ^30 11 12^.

Most significantly, SpatialCell AI achieves true single-cell resolution by directly measuring cell positions at pixel-level precision rather than computationally estimating them. This fundamental difference in approach enables the generation of individual cell records (Cell_1, Cell_2, Cell_3…) with precise coordinates and cell-specific expression profiles, rather than the enhanced spot-level data with cell type proportions produced by computational methods Table 1, Figure 6D.

### Performance and Clinical Translation Implications

The performance characteristics of these paradigms differ substantially. While computational methods show variable excellence depending on platform and reference quality—with top performers including CARD, Cell2location, Tangram, RCTD, and stereoscope—their performance is inherently constrained by the universal limitations described above. In contrast, SpatialCell AI maintains consistent high performance across all conditions due to its reference-independent, morphology-guided framework ^30 11 12^.

From a clinical translation perspective, computational methods face significant barriers due to their complex reference requirements, database dependencies, and platform-specific workflows. The need for matched scRNA-seq references and computational infrastructure for database management creates translation bottlenecks that limit clinical applicability. SpatialCell AI’s self-contained workflow, requiring no external databases or reference datasets, offers direct clinical applicability with standardizable protocols suitable for diagnostic applications.

## Discussion

The development of SpatialCell AI represents a fundamental methodological advance in spatial transcriptomics analysis, moving from computational inference to morphology-guided direct measurement. Our comprehensive validation demonstrates that this approach not only achieves single-cell resolution from spot-based data but does so with accuracy approaching native single-cell technologies (r=0.791), while eliminating the universal limitations that constrain all existing computational methods. The most significant contribution of SpatialCell AI lies in its significant departure from the expression-similarity paradigm that has dominated the field. Our analysis of 28+ computational methods across three major benchmarking studies ^30 11 12^ reveals that despite methodological diversity—spanning probabilistic models, matrix factorization, graph-based approaches, and deep learning—all share the same fundamental limitations: reference dependency, platform effects, sparse data sensitivity, and complete neglect of morphological information. SpatialCell AI addresses these limitations not through incremental improvements but through a methodological transformation that leverages the rich spatial and morphological context inherent in tissue architecture ^2, 3^. The implications of this methodological advance extend far beyond technical improvements. By achieving reference independence, SpatialCell AI enables analysis of novel tissues, rare diseases, and cross-species studies that were previously impossible with computational methods ^8, 9^. This capability is particularly crucial for precision medicine applications where patient-specific tissue analysis cannot rely on pre-existing reference databases.

Our distribution-based validation framework addresses a critical gap in the field—the lack of ground truth data for validating spatial-to-single-cell conversion methods. By using matched Xenium single-cell data as ground truth ^16^, we provide the first definitive evidence that computational enhancement can accurately reconstruct single-cell signals from multi-cellular measurements. The near-unity ratios for expression levels (0.97) and detection rates (1.02) after enhancement validate that SpatialCell AI recovers biologically accurate single-cell patterns rather than generating artificial signals ^20^. Particularly noteworthy is the framework’s consistent performance across different gene expression ranges. The high correlation for marker genes (r=0.870) ensures accurate cell type identification, while robust performance on lowly expressed genes (r=0.672) demonstrates sensitivity to subtle biological signals often lost in spot-based measurements. This dynamic range preservation is essential for detecting rare cell populations and understanding complex cellular states that define tissue microenvironments.

The self-contained nature of SpatialCell AI’s workflow presents new opportunities for clinical translation. Unlike computational methods requiring extensive reference databases and bioinformatic infrastructure ^11^,^12^, SpatialCell AI operates on standard histological images and spatial transcriptomics data available in routine clinical workflows. This simplicity, combined with standardized output formats compatible with existing analysis pipelines, positions SpatialCell AI as a bridge between research-grade spatial transcriptomics and clinical diagnostics. The framework’s ability to enhance lower-resolution, more affordable platforms (like standard Visium) to near single-cell resolution has profound implications for clinical accessibility. Rather than requiring expensive single-cell resolution platforms ^7^, clinical laboratories could achieve comparable results using established technologies enhanced by SpatialCell AI, democratizing access to high-resolution spatial biology.

While SpatialCell AI represents a significant advance, several limitations warrant consideration. The framework’s dependence on image quality for accurate cell segmentation may pose challenges in tissues with poor morphological preservation or low contrast. The computational requirements for processing large tissue sections, while manageable with modern infrastructure, may limit real-time applications. Future development should address these limitations through several avenues. The incorporation of multimodal data—such as protein expression from immunofluorescence—could further enhance cell type assignment accuracy ^2^. Additionally, developing lightweight versions optimized for edge computing could enable point-of-care applications.

## Conclusion

SpatialCell AI represents a significant advancement in spatial transcriptomics analysis, providing researchers with a powerful tool for cell-level investigation of tissue architecture. By integrating morphological and expression data, SpatialCell AI enables new resolution in spatial biology studies while maintaining compatibility with established analysis workflows.

The comprehensive validation using matched multi-platform data demonstrates that computational enhancement can accurately reconstruct single-cell expression patterns from spot-based measurements, achieving performance approaching true single-cell resolution technologies. The comparative analysis with existing computational methods reveals that SpatialCell AI not only addresses universal limitations but establishes an entirely new paradigm that transforms spatial transcriptomics from statistical inference to direct biological measurement.

This work establishes a new standard for validation in computational spatial transcriptomics and provides a robust framework for extracting single-cell information from widely-available spot-based platforms. The implications extend beyond methodological advancement to enable previously impossible applications in clinical diagnostics, novel tissue characterization, and high-precision spatial biology studies.

## Data and Code Availability

All validation was conducted using publicly available datasets to ensure reproducibility and eliminate potential bias. The complete validation framework including code for distribution-based comparison with Xenium ground truth is available at https://github.com/Spatialcell/spatialcell-ai-validation/tree/main. All SpatialCell AI-transformed single-cell resolution datasets generated in this study are freely available at https://zenodo.org/records/15794331. Raw spatial transcriptomics data from the original study ^16^ can be accessed through https://www.10xgenomics.com/platforms/visium/product-family/dataset-human-crc. Implementation details available through the corresponding author, with academic collaboration opportunities available upon request.

